# Mutational Signatures as Sensors of Environmental Exposures: Role of Smoking in COVID-19 Vulnerabilities

**DOI:** 10.1101/2021.09.27.461855

**Authors:** Yoo-Ah Kim, Ermin Hodzic, Ariella Saslafsky, Damian Wojtowicz, Bayarbaatar Amgalan, Teresa M. Przytycka

**Affiliations:** National Center for Biotechnology Information, National Library of Medicine, National Institutes of Health, Bethesda, MD 20894, USA

**Keywords:** SARS-CoV-2, mutational signatures, smoking, lung cancers, APOBEC, immune response to smoking, cell type composition, goblet cells, ciliated cells, basal cells

## Abstract

**Background:** Environmental exposures such as smoking are widely recognized risk factors in the emergence of lung diseases including lung cancer and acute respiratory distress syndrome (ARDS). However, the strength of environmental exposures is difficult to measure, making it challenging to understand their impacts. On the other hand, some COVID-19 patients develop ARDS in an unfavorable disease progression and smoking has been suggested as a potential risk factor among others. Yet initial studies on COVID-19 cases reported contradictory results on the effects of smoking on the disease – some suggest that smoking might have a protective effect against it while other studies report an increased risk. A better understanding of how the exposure to smoking and other environmental factors affect biological processes relevant to SARS-CoV-2 infection and unfavorable disease progression is needed.

**Approach:** In this study, we utilize mutational signatures associated with environmental factors as sensors of their exposure level. Many environmental factors including smoking are mutagenic and leave characteristic patterns of mutations, called mutational signatures, in affected genomes. We postulated that analyzing mutational signatures, combined with gene expression, can shed light on the impact of the mutagenic environmental factors to the biological processes. In particular, we utilized mutational signatures from lung adenocarcinoma (LUAD) data set collected in TCGA to investigate the role of environmental factors in COVID-19 vulnerabilities. Integrating mutational signatures with gene expression in normal tissues and using a pathway level analysis, we examined how the exposure to smoking and other mutagenic environmental factors affects the infectivity of the virus and disease progression.

**Results:** By delineating changes associated with smoking in pathway-level gene expression and cell type proportions, our study demonstrates that mutational signatures can be utilized to study the impact of exogenous mutagenic factors on them. Consistent with previous findings, our analysis showed that smoking mutational signature (SBS4) is associated with activation of cytokine-mediated signaling pathways, leading to inflammatory responses. Smoking related changes in cell composition were also observed, including the correlation of SBS4 with the expansion of goblet cells. On the other hand, increased basal cells and decreased ciliated cells in proportion were associated with the strength of a different mutational signature (SBS5), which is present abundantly but not exclusively in smokers. In addition, we found that smoking increases the expression levels of genes that are up-regulated in severe COVID-19 cases. Jointly, these results suggest an unfavorable impact of smoking on the disease progression and also provide novel findings on how smoking impacts biological processes in lung.

## 1 Background

Coronavirus disease (COVID-19) is an infectious disease caused by Severe Acute Respiratory Syndrome Coronavirus 2 (SARS-CoV-2). While most people infected with the SARS-CoV-2 virus experience mild to moderate respiratory illness and recover without requiring special treatment, some progress to the Acute Respiratory Distress Syndrome (ARDS). ARDS is a life-threatening, inflammatory lung injury that happens when fluid leaks into the lungs. There is a great heterogeneity within the population in susceptibility to the infection and to the progression of the disease to an acute stage. Complicating the analyses, the factors that facilitate infection of host organism by SARS-CoV-2 are not necessarily the same as the factors contributing to the progression of the disease to ARDS.

Cellular entry of the virus to host cells occurs through the binding of the viral spike (S) protein to host Angiotensin-Converting Enzyme 2 (ACE2) protein present on cell surface of many human cell types including respiratory and intestinal epithelial cells, endothelial cells, and kidney cells. The viral entry was also shown to require S protein priming and cleavage by TMPRSS2 protease [1].

In addition to ACE2 and TMPRSS2, several other genes have been suggested as factors facilitating the entry of the virus. Basigin (BSG) has been indicated as a potential alternative receptor [2] although the role has been debated in some studies [3]. Neuropilin-1 (NRP1) was also identified as a mediator of SARS-CoV-2 entry [4, 5]. Several proteases including cathepsin B/L (CTSB/CTSL) [6, 7], FURIN [8], TMPRSS4 [9] were considered as priming factors alternative to TMPRSS2.

Recent studies considered the impact of population level phenotypes as well as tissue and cell types on the expression of the SARS-CoV-2 entry proteins, focusing mainly on ACE2 and TMPRSS2 [10, 11, 12, 13]. Severity of COVID-19 has been associated with factors like age and sex on one side, and comorbidity such as hypertension, cardiovascular and respiratory diseases, cancer on the other [14]. Considering the wide spectrum of chemicals inhaled while smoking and known impacts of smoking on lung diseases including cancer, it is expected that smoking impacts both SARS-CoV-2 infection and COVID-19 progression. Initial studies focusing on the relation of smoking with ACE2 expression levels or on statistical analysis of disease cases based on smoking status produced mixed results. For example, it has been reported that tobacco smoking increases the expression of ACE2 in lung [15, 16] while other study showed that ACE2 content in the blood of healthy male volunteers was decreased by long term smoking experience [17] and even suggested that the risk of infection by COVID-19 might be reduced by half among current smokers [18]. Condition and tissue specific expression of ACE2 can, in part, contribute to these apparently contradictory results. In addition, factors contributing to the susceptibility to virus entry do not necessarily mirror the factors contributing to the likelihood of infected individuals to develop ARDS. For example, in the alveolar regions of the lung, ACE2 is expressed in type 2 pneumocytes (AT2 cells), which are fundamental for maintaining healthy tissue by secreting important surfactants and act as the progenitor cells for lung regeneration after injury [19]. Thus even if low ACE2 expression due to the reduction of AT2 cells could potentially decrease the opportunity of infection of aveoles, it might lead to a more severe outcome once infection happens.

Given the conflicting reports on the role of smoking for COVID-19, we turned to the large gene expression datasets of cancer and healthy tissues collected by the TCGA consortium in search for additional clues. In particular, we focused on the lung adenocarcinoma (LUAD) dataset since etiology of this cancer has many tangent points with COVID-19. This type of cancer develops typically in peripheral areas of the lungs and, as in the acute stage of COVID-19, shares similarities with pneumonia on a chest X-ray. Lung cancer patients often experience symptoms consistent with chronic obstructive pulmonary disorder (COPD) which predisposes them to ARDS. Most importantly, smoking is one of the key contributors to the emergence of LUAD and thus both cancer-free control and tumor samples can provide clues to the role that smoking has for the susceptibility to the SARS-CoV-2 infection and to the progression of COVID-19 to ARDS.

### Mutational signatures as sensors of environmental factors

Mutational signatures are characteristic mutation patterns imprinted on DNA molecules by specific mutagens [20, 21, 22, 23, 24, 25]. Such signatures can provide patient-specific information about environmental, immunologic, and other factors affecting the tissue of interest and allow us to decipher the impact of these factors on molecular pathways relevant to the disease of interest. For the technical details on the definition and inference of mutational signatures, we refer the reader to the pioneering paper of Alexandrov et al. [23] and a recent review [25]. Of importance for this study is that it is possible to infer whether or not a given signature associated with a specific mutagen has been active in the given genome as well as the strength of each signature, a value reflecting the exposure level of the given patient to the mutagen [25]. In particular, previous studies confirmed that the strength of the mutational signature related to smoking (referred to as SBS4) is indeed correlated with the exposure to smoking [21].

Mutagenic processes generating mutational signatures may be exogenous or endogenous. In particular, mutagenic processes that are caused by environment-related mutagens such as smoking are commonly considered as exogenous. In addition, for the purpose of this study, any other process that affects both tumor and healthy tissues of a patient simultaneously is considered to be exogenous. On the other hand, some endogenous processes such as dysfunction of DNA repair process is specific for tumor tissue and is often absent in healthy samples of the same individual. In this study, we focus on mutational signatures associated with environmental factors or with endogenous processes that affect both tumor and healthy tissues simultaneously and refer them as *exogenous mutational signatures*.

It is noteworthy that mutational signatures can be most easily inferred from sequencing of cancer samples. Since cancer is a result of clonal expansion, simple bulk sequencing can reliably capture somatic mutations that are common across cells in a cancer sample. In contrast, somatic mutations in healthy tissue are expected to occur randomly in different cells. Importantly, as healthy and cancer samples in the given tissue have been subjected to the same exogenous processes, signatures inferred based on cancer mutation data can be used to estimate the strength of exogenous factors, including smoking, acting on the healthy sample. Thus we postulate that the readouts of mutational signatures from cancer samples can be used to estimate the impact of exogenous processes on the tissue of interest (affecting both cancer and healthy samples). For gene expression, we utilize the data from normal (control) samples to examine changes in biological processes that are caused by the exogenous factors but are independent of cancer.

In this study, we investigated how genes and pathways with potential role for COVID-19 are related to exogenous factors such as smoking, using mutational signature exposures as measures of their strength (Fig. 1). In particular, by examining the genes whose expression profiles are correlated with the strength of smoking signature, we identified pathways which have a potential to impact the infectivity of the virus and progression to severe COVID-19 cases. In addition to changes in gene expression, we analysed changes in cell type composition caused by these factors. Overall, our results support the view that smoking is likely to contribute to the disease progression to an acute stage. Many of our findings are consistent with current knowledge, providing the confidence in our approach. Importantly, our analyses also revealed novel relationships that have not been appreciated previously, shedding more light on underlying biological processes associated with exposure to smoking and other exogenous factors.

**Figure 1:**
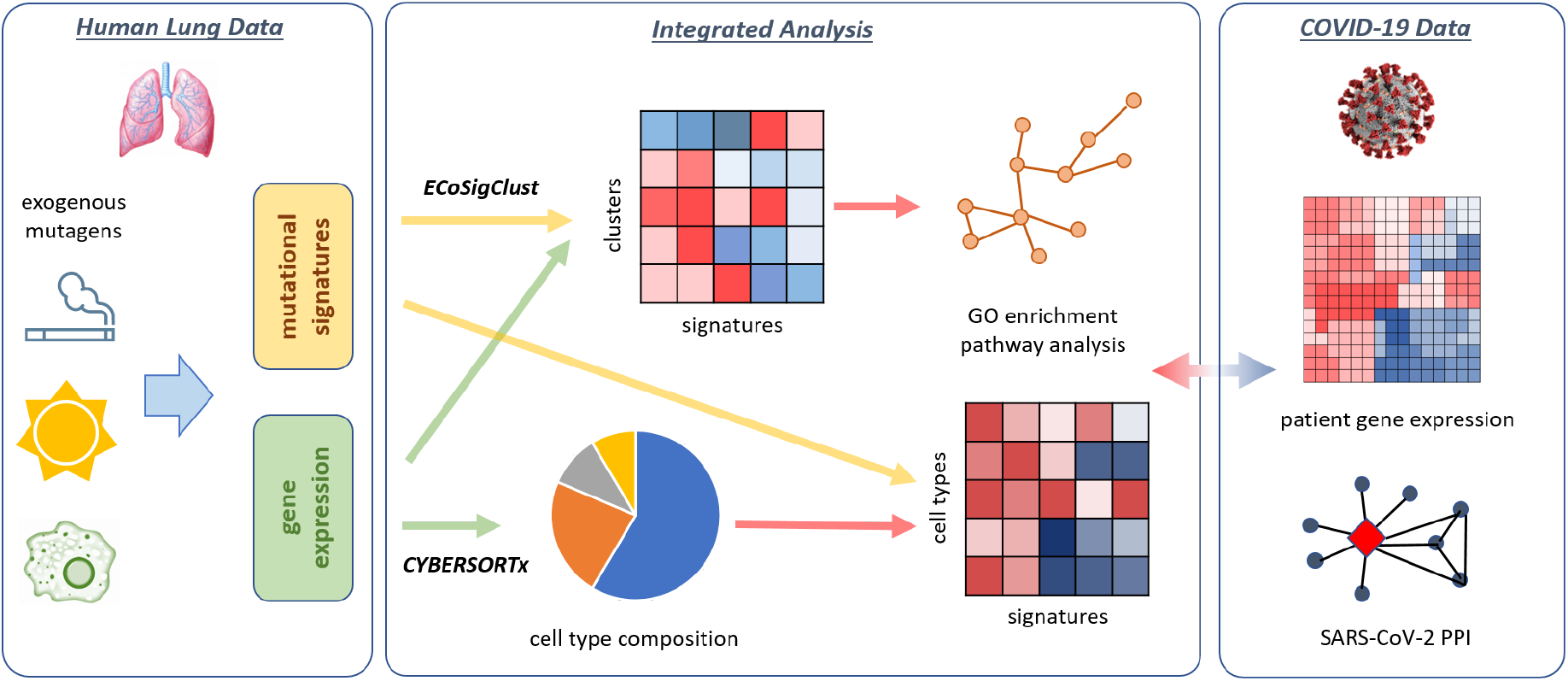
Overview of our integrated analysis of human lung and COVID-19 data. We utilize mutational signatures as sensors of exposure to exogenous mutagens. Combining with gene expression data, we uncovered biological processes associated with the exogenous mutagens and their potential impacts on COVID-19 susceptibilities. We also investigated changes in cell type composition caused by the mutagens.

## 2 Results

### 2.1 Mutational signatures in LUAD – Overview

We analyzed TCGA LUAD mutation data to infer mutational signatures in individual cancer patients using COSMIC signatures [20, 24] (See Methods). One of the primary mutagens contributing to the emergence of Lung Adenocarcinoma (LUAD) is smoking. Thus, genomes of LUAD patients harbor mutation patterns defined by smoking signature SBS4, which is assumed to reflect the level of the exposure to smoking. In addition to SBS4, the genomes of LUAD patients also harbor 5 additional COSMIC mutational signatures — SBS1, 2, 5, 13, and 40 (Methods).

Three of these signatures – SBS1, SBS5, and SBS40 – are often referred to as “clock-like” signatures as their strength is positively correlated with patients’ age in many (but not all) cancer types although differences in the etiology of the signatures are not fully understood. SBS5 is present in nearly all cancer types, and has been suggested to result from long-term or sporadic exposure to environmental causes such as oxidative stress [21, 26]. The accumulation of SBS40 mutations with age also suggests that it might be related to environmental factors and we found that it is strongly correlated with SBS4 in LUAD (Fig. S1). Finally, SBS1 is known to arise due to endogenous mutational process initiated by spontaneous or enzymatic deamination of 5-methylcytosine. SBS1 is not generally considered to be exogenous as favorable conditions for this process are during replication. Therefore, the signature exposures are likely to be different in tumor and normal samples due to the difference in replication rate.

The two remaining signatures, SBS2 and SBS13, are attributed to the mutations introduced by the AID/APOBEC family of cytidine deaminases enzymes, whose activity is generally related to immune response. Indeed, the strength of these signatures has been shown to be correlated with expression of immune genes and pathways [27]. It was not immediately obvious if SBS2 and SBS13 should be considered exogenous or not. The cause of the over-activity of APOBECs in LUAD are yet to be established, but Alexandrov et al. speculated that the cellular machinery underlying SBS2 and SBS13 can be activated by tobacco smoking, perhaps as smoking related inflammatory response [21], in which case, SBS2 and SBS13 would also be considered exogenous. Indeed, it is known that within the lung, cigarette smoke incites a potent inflammatory reaction in the airways and alveoli [28]. However, since we obtained the data from cancer patients, it is also possible that the immune response has been triggered by tumor cells. Nevertheless, it is interesting to understand the role of immune response independently of its triggers, which motivated us to include the APOBEC related signatures in our analysis.

### 2.2 SARS-CoV-2 entry gene expression levels are positively correlated with smoking related mutation counts

We leveraged the readouts of signature strengths to investigate if the factors causing mutational signatures, including smoking, can potentially impact the susceptibility to SARS-CoV-2 infection. To answer this question, we examined the correlation between the expression level of SARS-CoV-2 entry genes in normal tissue and the strength of mutational signatures. We first considered the established viral entry genes: ACE2, BSG, CTSB, CTSL, FURIN, NRP1, TMPRSS2, and TMPRSS4.

Focusing on the role of cigarette smoking, we found that the strength of SBS4 is positively correlated with the expression of most of these key genes facilitating the entry of SARS-CoV-2 (Fig. 2A). However, some of these correlations are not statistically significant, potentially due to the modest number of normal/control samples (*n* = 48). There is relatively strong correlation of smoking signature SBS4 with the expression of CTSB/L proteins as well as TMPRSS4. In addition, ACE2 has generally positive correlation with smoking signature although not statistically significant.

**Figure 2:**
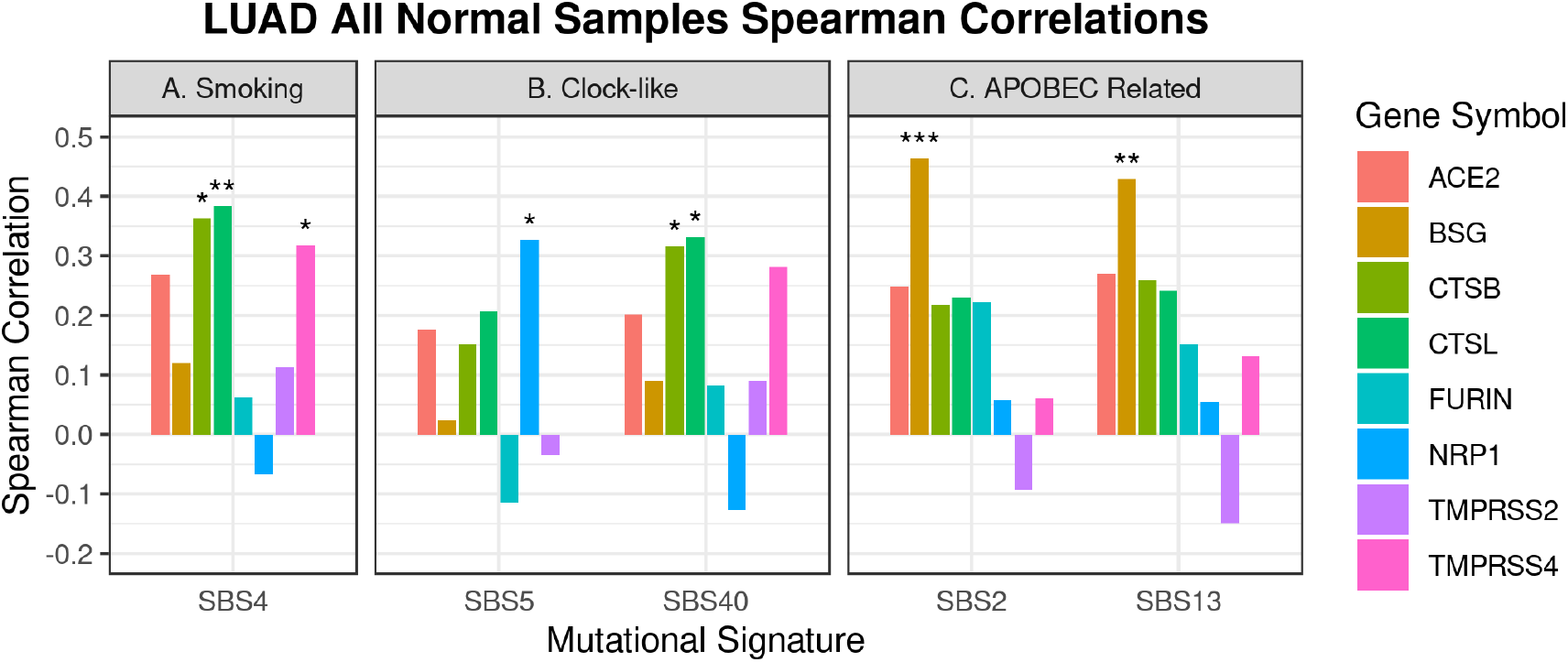
The correlation between mutational signatures and the expression of SARS-CoV-2 entry genes. The RNAseq data for TCGA LUAD normal samples are used while the mutational signatures are obtained from the matched cancer samples for the same patients. The pairs with statistically significant correlation are marked (\**p* ≤ 0.05, *\*\* p* ≤ 0.01, and *\*\*\* p* ≤ 0.001)

As for the two putatively exogenous clock-like signatures (SBS5 and SBS40), the correlations observed for SBS40 are similar to the ones for SBS4 while the activity of SBS5 has distinct patterns, indicating that different mutagenic processes are involved (Fig. 2B). SBS5 activities are positively correlated with NRP1. As mentioned above, it is assumed that NRP1 acts as a co-receptor for SARS-CoV-2 by binding furin-cleaved substrates and facilitates the infection of the virus. In addition, NRP1 is a cell surface receptor involved in the development of the cardiovascular system and angiogenesis [4, 5]. Finally, considering APOBEC related signatures, we found that the expression of BSG is significantly correlated with the strength of SBS2 and SBS13 (Fig. 2C). BSG (also known as CD147) is a member of the immunoglobulin superfamily. It is a multi-functional protein that plays a role in bacterial and viral infection as well as in cardiovascular diseases and thrombosis [29]. The former function provides its link to immune response while the later association is particularly interesting from the perspective of COVID-19. Thromboembolism risk of COVID-19 is believed to be high and associated with a higher risk of mortality [30]. Thus the prediction that BSG over-expression can be driven by immune system (possibly as an indirect consequence of cigarette smoking) is of primary interest. It is also interesting to note that BSG has been already proposed to be a promising target for COVID-19 treatment because of its potential involvement in an alternative route for SARS-CoV-2 invasion [31].

Overall our results suggest that smoking is not likely to reduce the probability of SARS-CoV-2 infection in lungs but instead may increase the likelihood of infection.

### 2.3 Pathway-based analysis of response to exogenous mutagenic processes

Having established that smoking has a potential to increase the risk of SARS-CoV-2 infection in lungs, we then asked how smoking can affect disease progression. Specifically, we wanted to learn how smoking and other exogenous mutagenic processes interact with cellular processes, in particular with processes known to be related to ARDS. Towards this end, we expanded the correlation analysis to all genes that are significantly correlated with at least one signature. Utilizing the approach developed in the previous study [27], we identified clusters of genes whose expression is correlated with different combinations of signatures (Fig. 3(a) and Table S1, Methods). More specifically, we selected the genes whose expression is significantly correlated with at least one mutational signature (*p* < 0.05) and clustered the genes based on their correlation patterns with mutational signatures. Many of SARS-CoV-2 entry genes belong to the clusters that are positively correlated with smoking signatures. We refer to this clustering procedure as `ECoSigClust` (**E**xpression **Co**rrelated **Sig**nature **Clust**ering).

**Figure 3:**
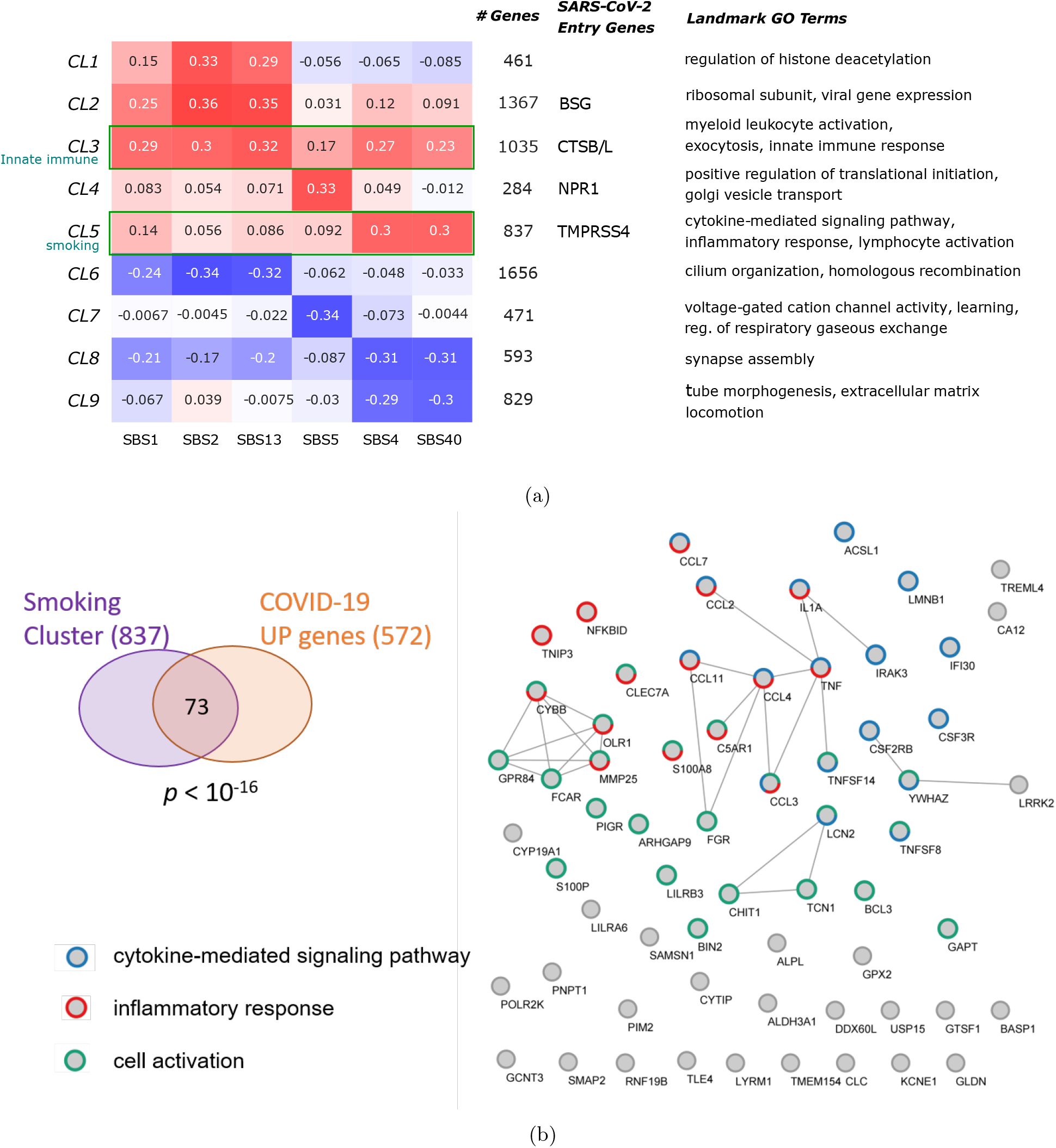
**(a)** `ECoSigClust` clusters based on the correlation between mutational signatures and gene expression. Genes having a significant correlation with at least one mutational signature (*p* < 0.05) are included in the clustering. The heatmap shows the mean correlation between signature and expression among all genes in the cluster (left). For each cluster, the number of genes, SARS-CoV-2 entry genes included in the cluster, and representative GO terms enriched in the cluster genes are also shown. **(b)** Smoking cluster genes overlapping with the genes upregulated in severe COVID-19 cases (COVID-19 UP genes). Interactions between genes (from STRING database) are shown if the combined interaction score is 0.8 or higher. The genes belonging to inflammatory response (red), cytokine-mediated signaling pathway (blue), cell activation (green) are marked.

GO Enrichment analysis of the clusters obtained by `ECoSigClust` revealed that the clusters are enriched with specific GO terms, providing insights on the interactions between signatures and molecular pathways. In particular, two clusters of genes have strong positive correlations with smoking signature – one cluster that is specifically correlated with SBS4 (Smoking cluster) and the other cluster which has overall positive correlation with all mutational signatures. Although both clusters include genes involved in immune responses, the genes in each cluster are associated with distinct mechanisms. In particular, the cluster with overall positive correlation with all signatures is enriched with the terms related to innate immune response, thus we refer to it as Innate immune cluster. Impacts of cigarette smoking on immune system has been appreciated before but not fully understood [32, 33]. Thus we analysed the two clusters further to obtain additional insights.

#### 2.3.1 The strength of smoking signature is correlated with increased inflammatory response

The cluster having the strongest positive correlation specifically with smoking-related signature (SBS4), which we call *smoking cluster*, includes 837 genes, enriched with cytokine-mediated signaling pathway (*p* < 10^*−*13^), inflammatory response (*p* < 10^*−*13^) and cell activation (*p* < 10^*−*14^). This is consistent with previous observations that the exposure of epithelial cells to smoking triggers pro-inflammatory response by altering signaling pathways such as mitogen-actived protein kinase (MAPK) or NF-*κ*B pathways and increases the release of pro-inflammatory cytokines and chemokines [32, 34], many of which are included in the cluster. Increased levels of pro-inflammatory cytokines and chemokines are associated with pulmonary inflammation in COVID-19 patients [35, 36], which points to potential relationship between inflammatory response to smoking and unfavorable progression of COVID-19.

To investigate whether the genes in the cluster are associated with severe cases of the disease, we compared each of the clusters with differentially expressed genes in post-mortem lung samples from COVID-19 patients [37]. Indeed, we found a significant overlap of the genes upregulated in the severe COVID-19 cases with the smoking cluster (73 out of 572 genes, *p* < 10^*−*16^, Fig. 3(b)) as well as another cluster (innate immune cluster discussed in Section 2.3.2) while no significant overlaps were found with other clusters. The overlapping genes include several chemokines (CCL2, CCL3, CCL4, CCL7, and CCL11), and pro-inflammatory cytokines (Interleukin 1*α* (IL1A), and tumor necrosis factor (TNF)). The significant overlap suggests that the cellular changes caused by smoking in host can contribute to the disease progression to the acute stage.

Interestingly, the smoking cluster includes MUC5AC gene, consistent with the previous report that cigarette smoking increases MUC5AC expression [38, 39]. MUC5AC is a canonical marker of mucus producing secretory goblet cells, which suggests the relation between smoking and goblet cells population as we confirm in Section 2.4. The overproduction of protective mucus is associated with the impairment of mucociliary clearance, an important first line defense mechanism in lung [40].

The cluster also includes TMPRSS4, an alternative protease promoting the entry of virus, which was found expressed in secretory cells of lung epithelium in previous studies [41, 42], supporting the expansion of secretory goblet cells caused by smoking. Another notable gene in the cluster is GPR15, a chemoattractant receptor for lymphocytes. The expression of GPR15 is found upregulated in smokers and known to be involved in T-cell trafficking [43].

Finally, the cluster also includes APOBEC3B, which is known to induce APOBEC related mutational signatures (SBS2 and SBS13). The fact that APOBEC3B belongs to the smoking cluster rather than the innate immune cluster supports the hypothesis that APOBEC enzyme over-activities generating the signatures are likely to be triggered by inflammatory response to smoking [21].

#### 2.3.2 Changes in innate immunity due to smoking may affect COVID-19 outcome

There is another cluster whose expression level is positively correlated with smoking signature SBS4 (Fig. 3(a)). The cluster, which we call *innate immune cluster*, includes 1,035 genes and is enriched with myeloid leukocyte activation (*p* < 10^*−*29^), secretion (*p* < 10^*−*19^), and intracellular transport (*p* < 10^*−*10^). Unlike the smoking cluster, this cluster is positively correlated across all mutational signatures including APOBEC-related signatures (SBS2 and SBS13). The exact mechanism triggering the over-activity of APBOECs is yet to be established but previous studies suggest that they are associated with smoking-related inflammation or response to foreign antigens (e.g., tumor neoantigens) [21, 44]. The fact that APOBEC3B belongs to the smoking cluster rather than the innate immune cluster supports the view that the APOBEC mutational signatures in LUAD are likely to be triggered by the response to smoking related-inflammation. Next, we observe that the innate immune cluster includes markers of myeloid leukocytes such as MRC1, CD14 (macrophages), S100A9 (neutrophils). This is consistent with previous studies indicating that exposure to cigarette smoking significantly affects the functions of innate immune systems by increasing the number of myeloid leukocytes [32]. The analysis of cell type composition discussed in Section 2.4 confirms and extends this finding. As with smoking cluster, this cluster has a significant overlap with upregulated genes in severe COVID-19 cases (86 out of 572 genes, *p* < 10^*−*18^, Fig. S2), suggesting further that smoking can predispose individuals to progression to an acute case of COVID-19.

Interestingly, the cluster includes two lysosomal cysteine proteases (CTSB/CTSL), both of which have been suggested to facilitate the entry of SARS-CoV-2 virus. In particular, the elevated expression of CTSL after COVID infection is reported and suggested as a promising target for anti-COVID drugs [45, 46, 47]. Given the importance of innate immune response in viral infection, we asked if this cluster includes genes recently found to be interacting with SARS-CoV-2 proteins [48]. Indeed, a large number of genes interacting with SARS-CoV-2 proteins (60 out of 389 genes, *p* < 10^*−*13^, Fig. S3) are also included in the cluster. For example, the cluster includes stress granule protein G3BP2 which Nucleocapsid (N) of SARS-CoV-2 virus binds to and CSNK2B, a subunit of CK2 linked to stress granule. Another example is members of CUL2 complex (CUL2 and ELOC) interacting with ORF10 protein. ORF10 may bind to the CUL2 complex and hijack it for ubiquitination [48]. The overlapping gene set is enriched with more specific immune processes such as cellular response to oxygen levels, protein transport, and neutrophil degranulation.

The innate immunity plays a critical role in dynamics of viral infection and disease progression [49], and smoking-related innate immune responses, together with significant overlap with COVID-related genes suggest that smoking may negatively impacts COVID-19 outcome.

#### 2.3.3 Additional insights from other smoking related ECoSigClust clusters

While we did not find significant overlaps of other clusters with differentially expressed genes in severe COVID-19 cases reported in Blanco-Melo et al. [37], GO enrichment analysis for these clusters provided interesting insights on the impact of smoking on cellular function and further demonstrate the power of our approach using mutational signatures. For example, we observe that smoking signature SBS4 is negatively correlated with Clusters 8 and 9, both of which are enriched with cell differentiation and morphogenesis.

SBS5 is correlated negatively with Cluster 7, which is enriched with genes related to voltage gated cation channel activity and neurotransmitter receptor complex. It is known that these channels are targets of a number of naturally occurring toxins, therapeutic agents as well as environmental toxicants [50], including nicotine [51]. In addition, ion channels are key components of cilia signalling. Importantly, the cluster also contains known early transcriptional drivers of ciliogenesis such as MYB and TP73, consistent with the reports that smoking blocks early ciliogenesis [52, 53]. This suggests that reduction in expression of genes associated with ion channels might be related to the reduction of ciliated cells population. The results discussed in Section 2.4 provide further insights into this relation.

In addition to the positive correlation with Cluster 3, the two APOBEC signatures (SBS2 and SBS13) are positively correlated with the expression of genes in Cluster 1 and 2, and negatively correlated with Cluster 6. In particular, Cluster 2 includes BSG, a gene indicated in SARS-CoV-2 entry discussed in Section 2.2. This suggests that inflammatory response to smoking might induce higher expression of BSG and contribute to COVID-19 infectivity. In addition, a recent study linked the expression of BSG gene to increased mucus secretion induced by cigarette smoke in chronic obstructive pulmonary disease (COPD) [54].

Finally, Cluster 6 includes TUBB1, a marker of ciliated cells. Interestingly, GO enrichment analysis of Cluster 6 found that this cluster is significantly enriched with cilium, linking the decreased proportion of ciliated cells with APOBEC activity and immune response. We subsequently confirmed this relation by the cell composition analysis reported in the next subsection.

### 2.4 Mutational signatures reveal relation between the exposure to exogenous processes and cell type composition changes

The signature dependent expression changes of MUC5AC, a canonical marker of mucus producing secretory goblet cells, as well as other markers of myeloid leukocytes discussed in the previous section suggest changes in the cell type composition associated with mutational signatures. Indeed, previous studies reported that exposure to smoking leads to the expansion of mucus secreting goblet cells and the decreased number of ciliary cells [41]. Other studies reported that cigarette smoke exposure reduces the number of ciliated cells and leads to functional and structural abnormality of the cells [55]. To investigate the correlation between cell type composition and mutational signatures, we decomposed the bulk expression data using CIBERSORTx and estimated cell composition in each sample (See Methods). Considering epithelial and immune cells separately, we then computed the correlation coefficients between the proportion of cell types (within epithelial and immune cell types, respectively) and the strength of mutational signatures (Fig. 4), which revealed several changes in both epithelial and immune cell type composition correlated with the mutational signature activities.

**Figure 4:**
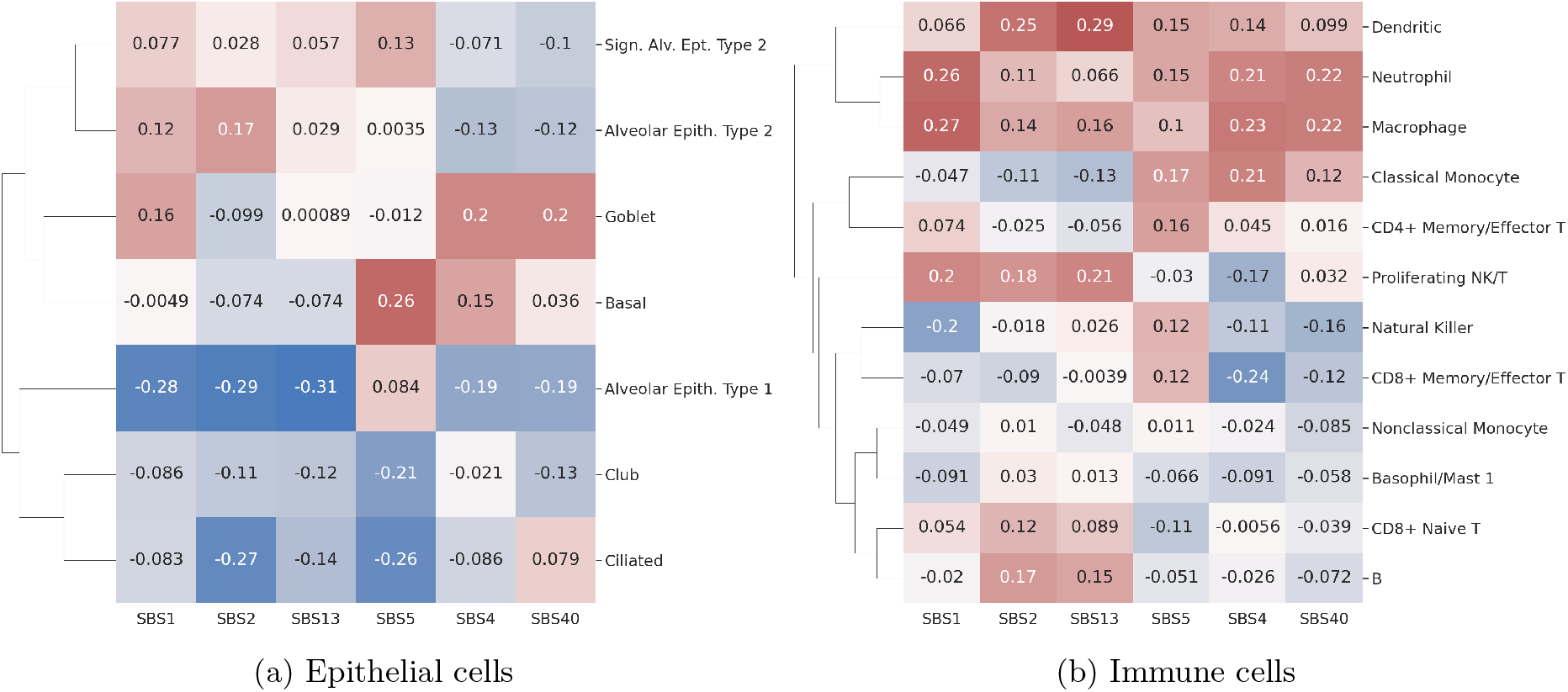
Correlation between mutational signatures and cell composition. Bulk expression counts are decomposed into different cell types using CIBERSORTx, and Spearman correlation coefficients are shown for (a) epithelial cells and (b) immune cells, separately.

Among epithelial cell types, the proportion of goblet cells and to a lesser extend basal cells is positively correlated with smoking signature SBS4 and all other cells have negative or no correlations (Fig. 4(a)), which confirms the previous observation that the exposure to cigarette smoke increases the number of mucous secreting goblet cells and thus can lead to goblet cell hyperplasia, mucus hypersecretion and promote inflammatory responses. The correlation pattern of goblet cells is similar to the pattern of smoking cluster in Fig. 3(a) (positive correlation with SBS4 and SBS40, and modest positive correlations with SBS1), supporting the hypothesis that the inflammatory responses are generated by epithelial cells with altered cytokine-mediated signaling pathways in response to smoking exposure. A previous study found that bronchial epithelial cells exposed to cigarette smoke produced a dose-dependent increase in the expression of MUC5AC, IL8 (also called CXCL8) and TNF*α* genes [56], all of which belong to the smoking cluster. Basal cells are located below the surface epithelial cell layer and serve as progenitor cells from which ciliated, secretory, and goblet cells differentiate during repair. While the increases of basal cells in smokers has been suggested before [41], it is interesting to note that the proportion of basal cells correlate most strongly with SBS5.

The proportion of ciliated cells has negative correlations with SBS2 and SBS5. The major function of airway ciliated cells is to mediate propulsion of mucus gel. Thus a proper balance between goblet and ciliated cells is required for correct functioning of lungs. Previous studies indicated that this balance might be disturbed by smoking [41, 55]. For example, cigarette smoke exposure is associated with reduced number of ciliated cells as well as with functional and structural abnormality of the cells [55]. Our results additionally reveal that the decrease in the number of ciliated cells is most strongly related to SBS2 and SBS5. Indeed, in addition to SBS4, mutation landscape of smokers is characterised by increased activities of SBS5 [21]. Although the causes of SBS5 are not fully understood, some studies hypothesize that SBS5 may, in part, emerge due to a long term exposure to chemicals including nicotine [57]. In addition, the reduction of the number of ciliated cells is associated with SBS2, suggesting a potential relation to immune response.

Finally, of particular interest is the dynamics of the population of AT2 cells. A recent study demonstrated that AT2 cells are significantly decreased in COVID-19 patients [58]. Our results show that the population of AT2 cells decreases also with smoking. In contrast, the proportion of signaling AT2 cells, a subtype of AT2 cells that can serve as progenitor cells for restoring population of alveolar cells [59] increases with SBS5. This observation is consistent with previous studies suggesting that exposure to cigarette smoke enhances the stemness of AT2 cells [60]. It is interesting that the proportion of signaling AT2 cells positively correlates with SBS5, a signature that accompanies smoking.

As for immune cells, we observe that innate immune cells such as dentritic cells, macrophages, and neutrophiles have overall positive correlation across all mutational signatures (Fig. 4(b)). This pattern is consistent with the positive correlation pattern of the innate immune cluster obtained with `ECoSigClust` (Fig. 3(a)), confirming changes in innate immunity due to cigarette smoking exposure.

In summary, the decomposition of expression into different cell types supports the inflammatory responses of epithelial cells and the increased innate immune activities with the exposure to smoking. The significant overlap with the inflammatory and innate immune response genes with differentially expressed genes in severe COVID-19 cases suggest that the tissue damage and impaired innate immunity due to smoking may negatively affect the disease progression. We also observe a reprogramming of epithelial cell composition associated with smoking that parallels the changes related to COVID-19 disease.

## 3 Conclusions

Although it has been appreciated that the exposure of individuals to environmental factors such as smoking or various chemicals might influence vulnerability to diseases, the level of such exposure has been difficult to quantify. Interestingly, many of such adverse environmental factors are mutagenic and leave characteristic mutational signatures in the genome. With the availability of large cancer data sets, computational methods have been developed to identify a number of mutational signatures, some of which are related to environmental factors. Most of analyses of mutational signatures have been focused on the role of mutagens in cancer. However, our study shows that the utility of mutational signatures can go beyond cancer studies and help shed light on the role of environmental mutagens as risk factors of other diseases.

Currently, mutational signatures are most readily accessible for cancer patients. We reasoned that even if the signatures are inferred from mutations in cancer cells, exogenous environmental factors equally impact cancer and non-cancer cells. Therefore, in this study, we used signatures obtained from cancer genomes to infer the exposure level to environmental factors and gene expression from control tissues to study the activity of molecular pathways related to the mutagenic environmental processes. One should keep in mind that only the signatures of exogenous processes acting on both cancer and healthy tissue can be used for this approach. While it is clear that smoking is an exogenous mutagen, the etiology of many signatures is still unknown and it might be difficult to confidently establish whether a given signature should be considered as exogenous or not. To avoid this uncertainty, it is thus desirable to include DNA sequencing in future single cell studies in addition to expression measurements.

In addition to utilizing exogenous signatures, it was important to use expression data from non-cancer tissues as the impact of mutagens on gene expression might be quite different in cancer and in healthy tissue. Indeed, different from normal samples, the expression of most of COVID-19 entry genes have negative correlation with the smoking signature in cancer samples (Fig. S4). Although the expression of CTSL remains positively correlated with SBS4, the correlation is less significant in cancer samples compared to normal samples. The drastic difference may be due to the changes in the proportion of cell types in cancer samples.

Our signature-based analysis uncovered many interesting insights on how smoking can impact activities of genes and pathways related to COVID-19. The results of our studies are in good agreement with current knowledge, adding confidence in the approach. Furthermore, our results provide additional insights that were not accessible with previous approaches. For example, previous studies suggested smoking can decrease ciliated cells and increase goblet cells in their proportion [41, 61]. By utilizing continuous mutational signature values rather than binary smoking status, our analysis further revealed that the decrease of ciliated cells is, in fact, related to the activity of the SBS5 signature that accompanies smoking. Of importance to COVID-19 is the observation that smoking leads to reduction of proportion in alveolar cell type 1 progenitor (AT1) cells. This reduction was implicated by the results of CIBERSORTx. AT1 cells enable breathing and gas transfer and therefore, their reduction might result in breathing impairments in heavy smokers, and likely to predispose to breathing problems in COVID-19 patients. The interplay between smoking and immune system that we uncovered was also consistent with current knowledge although the concern is still remaining that some of the immune response in the control tissue could be contributed by immune response to cancer. Finally, significant overlap of upregulated genes in severe COVID-19 cases with smoking and innate immune clusters as well as the enrichment of interactors with SARS-CoV-2 proteins in innate immune cluster provide additional evidence for the potential impact of smoking on COVID-19 infection and progression.

Overall, our approaches of looking at the data through the lenses of mutational signatures provide a new and powerful stepping stone for studying the potential impact of environmental factors on individual’s health, disease susceptibility and progression. We have demonstrated the utility of our approach in the context of lung diseases by showing that the relationship between mutational signatures and gene expression are consistent with current knowledge, and by uncovering additional underlying dependencies that shed light on the relation between COVID-19 and smoking.

## 4 Methods

### Mutational signatures

We downloaded the TCGA LUAD (lung adenocarcinoma) exome mutation spectra (528 samples) and the exome COSMIC reference mutational signatures, provided by Alexandrov et al. [24], from Synapse (accession numbers: syn11801889 and syn11726602, respectively). To determine the predominant signatures being active in LUAD samples, we started with the initial sample exposures to mutational signatures from [24] (Synapse accession number: syn11804065). The list of active signatures was refined to remove any rare signatures; namely we keep only signatures that were present in at least 5% of samples and were responsible for at least 1% of mutations. Next, using such a list of active mutational signatures in LUAD (SBS1, SBS2, SBS4, SBS5, SBS13, SBS40, and SBS45), we determined their sample-specific exposures using the quadratic programming (QP) approach available in the R package – SignatureEstimation [62]. Signature SBS45 was omitted from the analyses presented in this study as this signature is likely an artefact due to 8-oxo-guanine introduced during sequencing (see COSMIC Mutational Signatures website: https://cancer.sanger.ac.uk/signatures/).

### Expression data

TCGA LUAD RNAseq expression data were obtained from the Genomic Data Commons Data Portal (https://portal.gdc.cancer.gov/) on June 5th, 2020. HTseq counts were normalized and variance-stabilizing transformed (vst) using DESeq2 [63]. Only donors that had both gene expression and mutational signature exposures were kept, which resulted in 48 normal samples and 466 tumor samples used in this study.

### Clustering

To identify expression based pathways that are associated with signatures, we used `ECoSigClust` developed for our previous analysis [27]. Specifically, we first computed Spearman correlation coefficients of the expression level and mutation counts for each pair of genes and mutational signatures. We then selected the genes exhibiting significant correlation with at least one of the mutational signatures; the expression of a gene is considered significantly correlated with a signature if nominal *p* < 0.05. This procedure selected 7,533 genes. We then clustered the genes based on their correlation patterns using a consensus K-means algorithm; running K-means clustering 100 times with random start and varying *k* from 5 to 50 and subsequently running hierarchical clustering with the consensus matrix from 100 runs of K-means algorithm. To determine the optimal cluster number, three different clustering validation metrics – Silhouette Index, Calinski-Harabasz Index, and Davies-Bouldin Index – were used, measuring compactness within clusters and separation between clusters slightly differently. The chosen number of clusters *k* = 9 was based on these metrics (Fig. S5) and was kept small for interpretability of each cluster. GO enrichment analysis was performed using hypergeometric test for each cluster with all genes included in the clustering as background to assess the differences among the clusters. The list of genes and enrichment analysis results for all clusters are provided in Table S1.

### Differentially expressed genes in patients with severe COVID-19

To evaluate how the genes in the identified clusters are affected in patients with severe COVID-19 cases, we compared the genes in each cluster with differentially expressed genes in post-mortem lung samples from COVID-19-positive patients (Supplementary Table S4 in [37]). We considered the genes with positive (resp. negative) fold change and adjusted *p*-value < 0.05 as significantly upregulated (resp. downregulated).

### Interactors with SARS-CoV-2 proteins

To interrogate whether the genes in the identified clusters encode proteins interacting with the SARS-CoV-2 proteins, we utilized the interactome data collected by Gordon et al. [48]. High-confidence interactors with SARS-CoV-2 proteins compiled in Supplementary Table S2 from the study is included.

### Cell composition analysis with CIBERSORTx

HTseq raw counts in bulk expression data for the normal samples from TCGA LUAD dataset were used for the analysis. For each gene, the counts in every sample were normalized by the total sum of counts in that sample, multiplied by 1 000 000. The genes without at least one normalized count with a value greater than 1 were discarded. The *Human Lung Cell Atlas* (HLCA) [64] single cell reference data containing 42 distinct cell types was obtained in form of counts from Synapse (accession number: syn21560511). As per CIBERSORTx guidelines, the same normalization procedure was used on the single cell reference data and used as an input to CIBERSORTx to impute the cell proportions of the 42 given cell types in the bulk TCGA-Lung expression data.

For two subsets of cell types - epithelial and immune cell types, we computed Spearman correlation of each imputed cell type’s fraction with the exposures of Signatures 1, 2, 4, 5, 13, and 40. The strength of the correlation and the resulting heatmaps are shown in Fig. 4.

## Acknowledgements

This research was supported by the Intramural Research Program of the National Library of Medicine at the NIH.

## Availability of data and materials

No data or materials are generated in this study. `ECoSigClust` is available at https://github.com/ncbi/ECoSigClust.

## Supplemental Information

**Figure S1:**
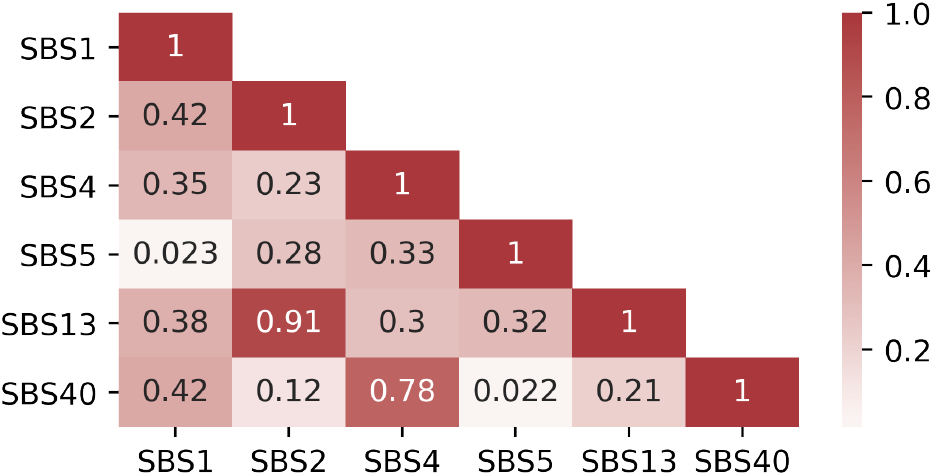
The Spearman correlation coefficients among the mutational signatures in TCGA LUAD for the patients with RNAseq data from normal samples used in this study.

**Figure S2:**
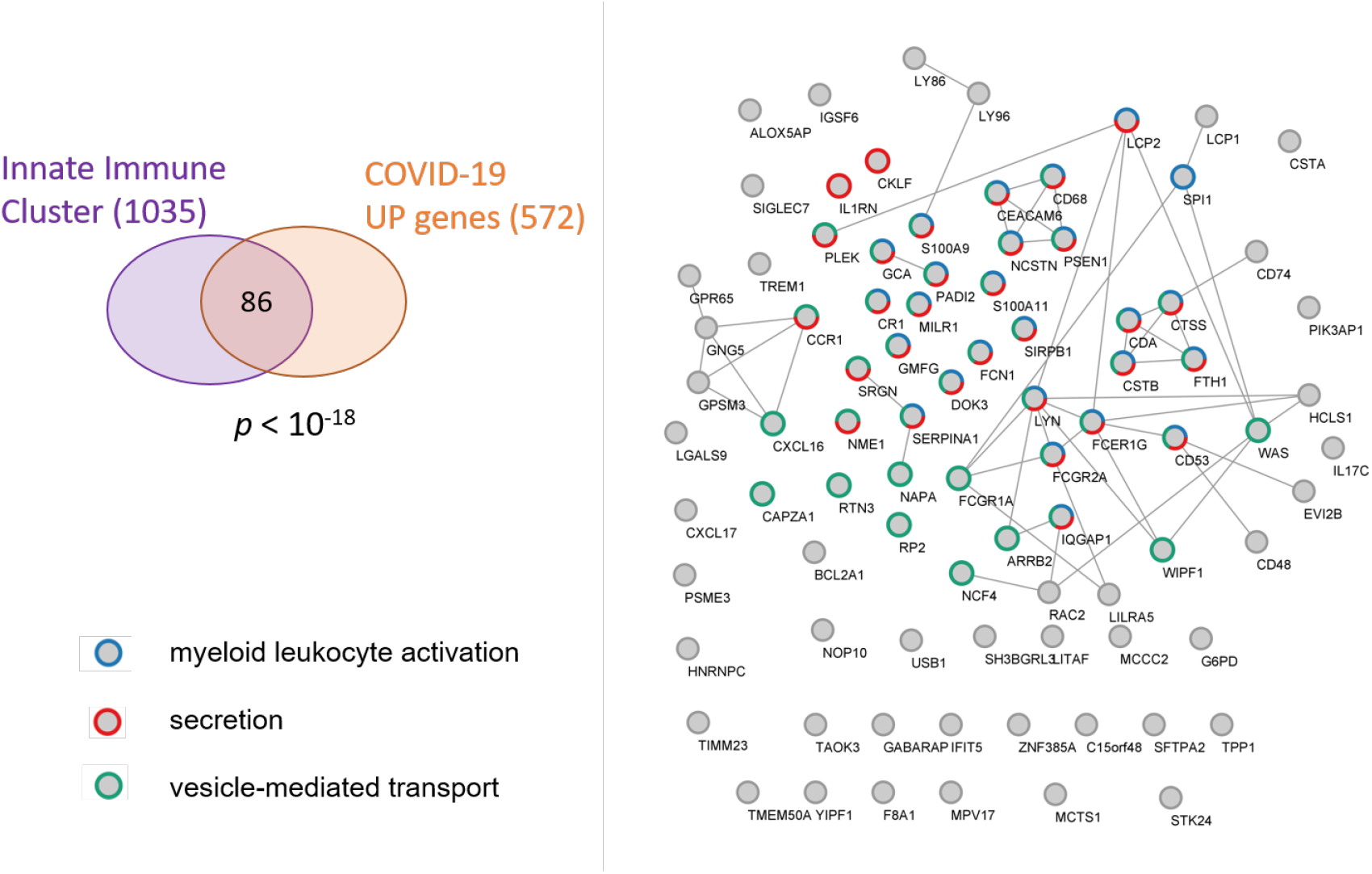
Innate immune cluster genes overlapping with the genes upregulated in severe COVID-19 cases. The interactions with combined score of 0.8 or higher in STRING database are shown. The genes belonging to myeloid leukocyte activation (blue), secretion (red), and vesicle-mediated transport (green) are marked.

**Figure S3:**
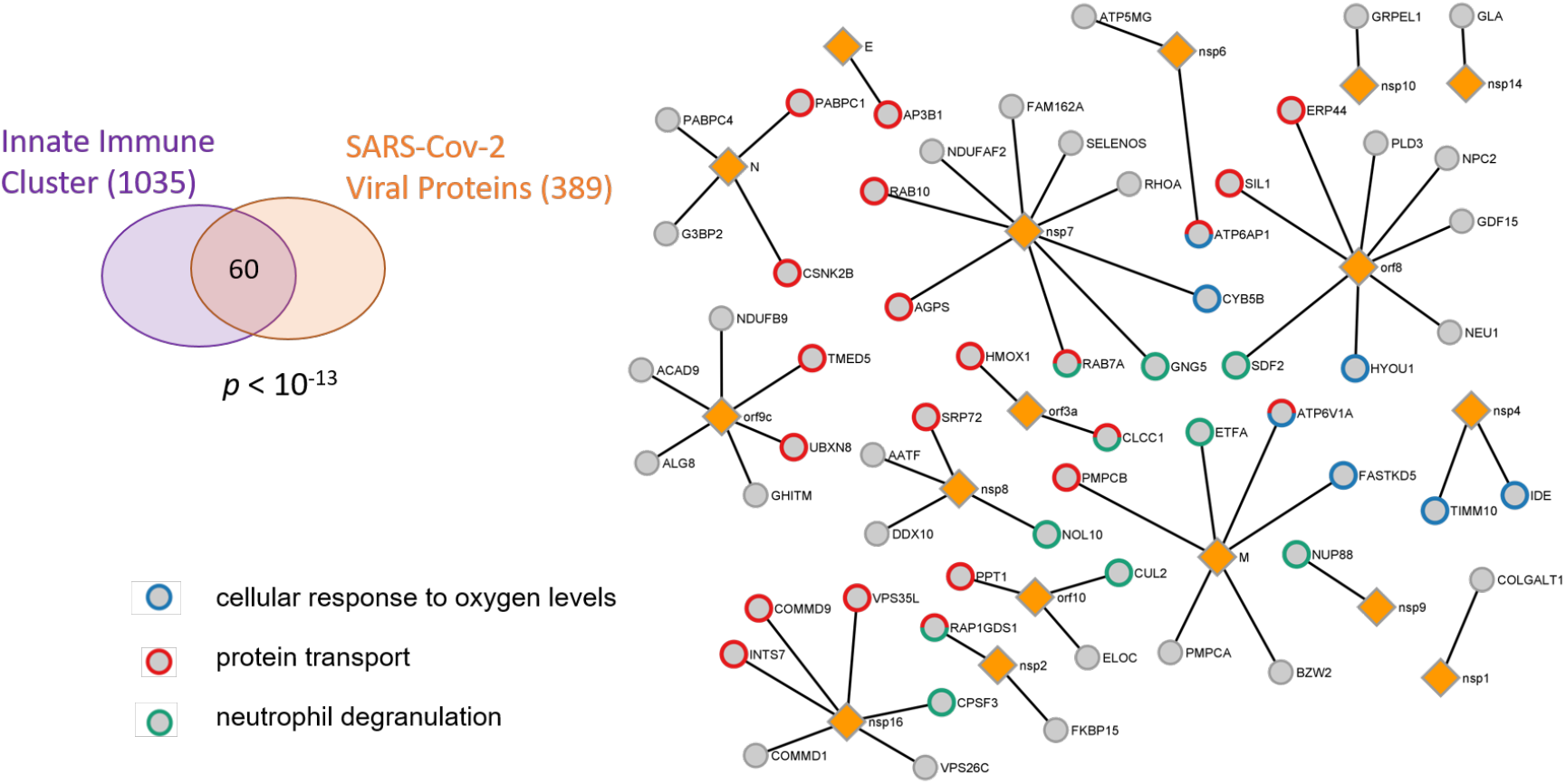
Innate immune cluster genes (circles) interacting with SARS-CoV-2 proteins (orange diamonds). Viral proteins interacting with each gene is shown. The human genes belonging to cellular response to oxygen levels (blue), protein transport (red), and neutrophil degranulation (green) are marked.

**Figure S4:**
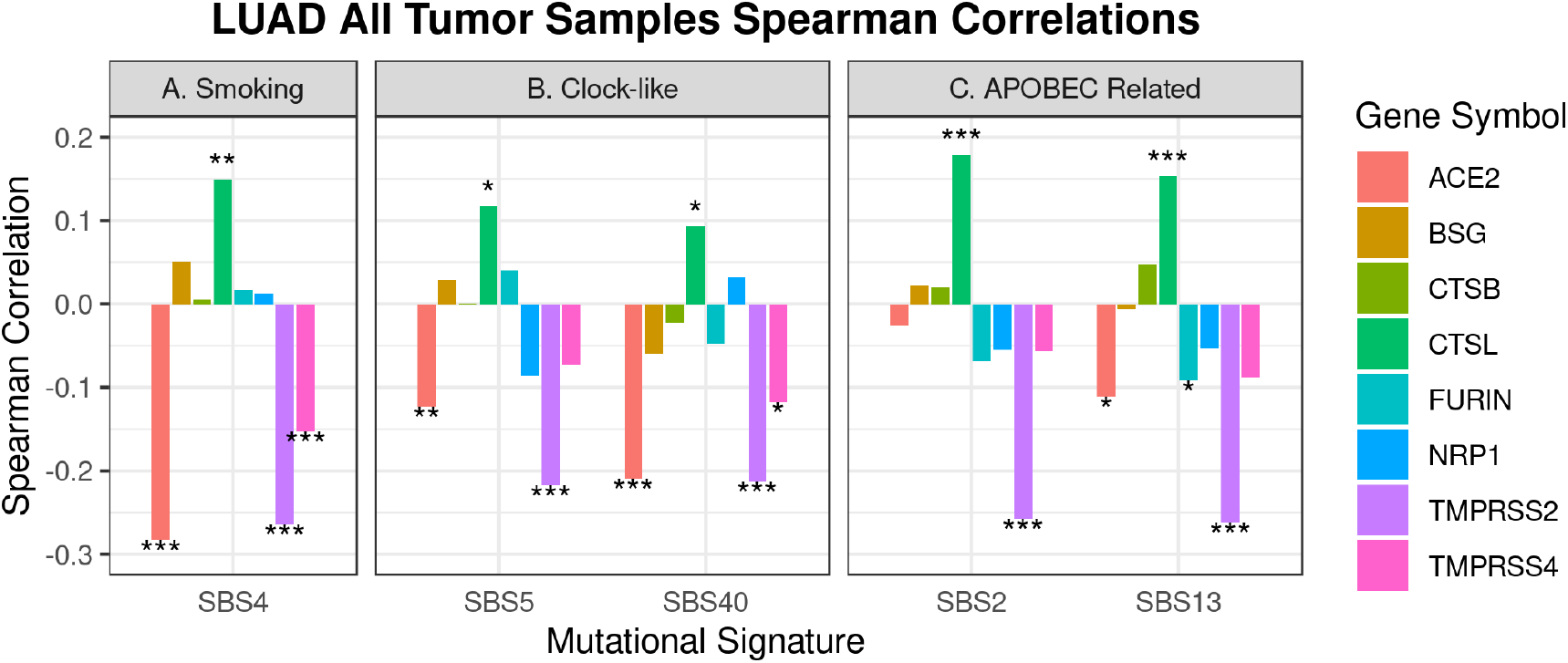
The correlation between mutational signatures and the expression of SARS-CoV-2 entry genes. The RNAseq data and the mutational signatures for TCGA LUAD tumor samples is used. The pairs with statistically significant correlation are marked (\**p* ≤ 0.05, *\*\* p* ≤ 0.01, and *\*\*\* p* ≤ 0.001)

**Figure S5:**
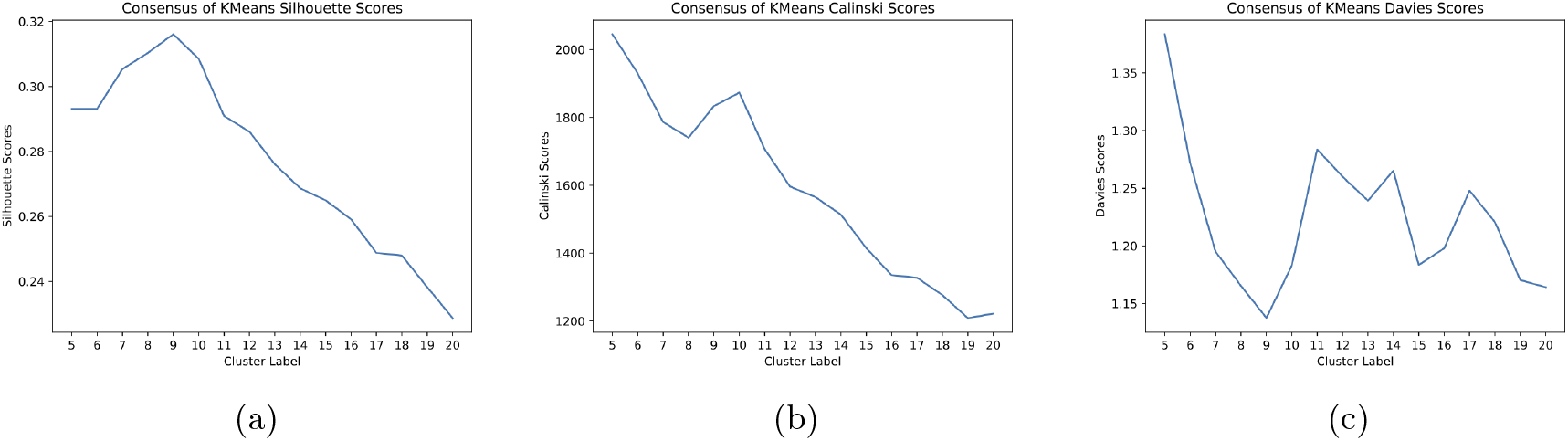
Evaluation of clustering for varying *k*’s (the number of clusters) using different metrics: (a) Silhouette Index, (b) Calinski-Harabasz Index, and (c) Davies-Bouldin Index.

## Notes

### Competing Interest Statement

The authors have declared no competing interest.

